# ADOLESCENT ALCOHOL EXPOSURE DISRUPTS ASTROCYTE-SYNAPTIC STRUCTURAL AND FUNCTIONAL COUPLING IN THE MALE DORSAL HIPPOCAMPUS

**DOI:** 10.1101/2025.10.14.682478

**Authors:** O. Coulter, C.D. Walker, T. Carter, H.G. Sexton, J. Denvir, W.C. Risher, B.J. Henderson, M-L Risher

## Abstract

Adolescence is a window of heightened vulnerability to the neurotoxic effects of binge ethanol exposure. Adolescent intermittent ethanol (AIE) exposure has been shown to induce long-lasting cognitive and behavioral impairments in patients and rodent models that increase the risk of developing alcohol use disorder (AUD). Our previous work shows that these behavioral deficits coincide with persistent astrocyte dysfunction. Here, we aim to understand how astrocyte-synaptic structural and functional crosstalk are disrupted following AIE to provide better mechanistic understanding of why behavioral impairments persist into adulthood. Male Sprague-Dawley rats received AIE, a variety of adeno-associated viruses encoding astrocyte-specific sensors, and fiber implantation in the dorsal hippocampal (dHipp) for *in vivo* photometry. A subset of rats received hM3D(Gq) to chemogenetically activate astrocytes. Following AIE and a forced abstinence period that allowed growth into adulthood, rats underwent assessment in the contextual fear conditioning (CFC) task with simultaneous fiber photometry recordings. By combining immunohistochemistry (IHC), Stimulated Emission Depletion (STED) microscopy, fiber photometry, chemogenetics, and slice physiology, we show that AIE induces structural and functional decoupling of astrocytes from synapses and astrocyte dysregulation that persists into adulthood. Remarkably, stimulating astrocytic calcium signaling via chemogenetic activation partially attenuates heightened fear responding and increases gliotransmitter availability. These findings highlight a critical role for astrocyte-synaptic crosstalk in regulating fear learning and underscore the untapped therapeutic potential of targeting astrocytes to improve behavioral outcomes following substance use.

## Introduction

In the United States, alcohol use typically begins during adolescence with consumption escalating throughout the early-to-mid-twenties. Approximately 90% of adolescent consumption is considered binge drinking (4 drinks for a female and 5 drinks for a male within a 2-hr period) with over 35% reporting regular consumption^1^. Binge-like consumption during adolescence has been associated with fear behavior dysregulation^2,3,^ that can worsen mental health outcomes and contribute to the onset of excessive use of illicit drugs. Equally problematic are the long-term consequences that can include chronic health issues, the emergence of psychiatric disorders, persistent cognitive deficits, and increased susecptibility to alcohol and substance misuse^4,5,6^. It has been proposed that this is, in part, due to the convergence of binge drinking with ongoing brain development, resulting in maladaptive brain maturation that drives permanent alterations in neuronal function and connectivity^6^.

With rodent models of adolescent intermittent ethanol (AIE) exposure, there is growing evidence demonstrating that adolescent binge drinking compromises neuronal function and associated cognitive and emotional regulation^7-11^. Non-neuronal cells, particularly astrocytes, are key modulators for these outcomes by regulating synaptic structure, function, and cognitive behavior^12-15^. Astrocytes are specialized glial cells that work extensively within the central nervous system (CNS) to maintain the extracellular environment^16^, provide neuroprotection^17^, and influence synaptic transmission^18,19^. Bidirectional communication between astrocytes and synapses is facilitated via the ensheathment of pre- and postsynaptic terminals by peripheral astrocyte processes (PAPs). Appropriate PAP ensheathment of excitatory synapses is critical for maintaining an optimal homeostatic environment. PAP-synaptic proximity has been shown to result in a number of changes within the tripartite synapse that are critical for their optimal function^20-22^. Despite our growing awareness of how this interaction maintains synaptic health and function, the specific role of astrocytes in AIE-induced changes in neuronal function and behavior remain unclear.

Previous work from our lab and others has demonstrated that AIE results in a reduced threshold for long-term potentiation (LTP) and excessive acute and chronic neuronal excitability that persists into adulthood, despite a multiple weeks of forced abstinence^20,23^. The persistent increase in neuronal excitability correlates with the chronic upregulation and release of astrocyte secreted factors that are known to interact with neuronal receptors, influencing synaptogenesis and dendritic spine maturity^24^. Interestingly, the expression of astrocyte secreted factors increases during abstinence and coincides with a shift towards a more immature dendritic spine phenotype within the CA1 dorsal hippocampus (dHipp)^20^. Previous work using 3D electron microscopy to reconstruct male adult rat dHipp shows that reduced PAP ensheathment and proximity to local synapses correlates directly to the level of dendritic spine maturation in the adult dHipp, with more immature spines having reduced PAP coverage^25^. These findings, combined with a growing appreciation for the role that astrocytes play in synaptic maturation, regulation, and behavioral outcomes, provide the rationale for investigating the long-term effects of AIE on astrocyte morphology, PAP-synaptic proximity, and neuronal-astrocyte communication in the adult male CA1 dHipp. We hypothesize that the loss of dendritic spine maturity is due to an AIE-induced loss of PAP-synaptic proximity, driving impaired neuronal-astrocyte communication and contributing to the dysregulation of neuronal activity and behavior. We further hypothesize that activation of dHipp astrocyte-targeted Designer Receptors Exclusively Activated by Designer Drugs (DREADDs) is sufficient to rescue AIE-induced disruption of hippocampal-dependent behavior and gliotransmission during a contextual fear conditioning task.

## METHODS

### Animals

All procedures in this study were conducted in accordance with the guidelines of the American Association for the Accreditation of Laboratory Animal Care and the National Research Council’s Guide for Care and Use of Laboratory Animals and were approved by the Institutional Animal Care and Use Committee at Marshall University and the Huntington Veterans Affairs Medical Center. 178 male Sprague Dawley rats (Hilltop Lab Animals Inc., Scottdale, PA) were received on postnatal day (PND) 24. Animals were double housed with cagemates and randomly assigned to experimental groups prior to EtOH exposure, with experimenters blinded throughout all experiments and data collection. Analyses were conducted with experimenters blinded to the assigned experimental groups. All animal numbers were determined based on the individual experimental group sizes outlined below and in a prior power analysis using G*Power version 3.1.9.4 (Axel Buchner, Heinrich Heine University, Düsseldorf, Germany) for sample size estimation. This analysis was informed by previously published data and pilot findings, using a significance threshold of α = 0.05 and power β = 0.8 to determine the minimum required sample size.

### Intermittent Binge EtOH Exposure

Animals received ethanol (EtOH; cat #89125-162; Koptec 190 Proof Pure Ethanol; Avantor; Radnor, PA) or water (H_2_O) as previously described^26^. Beginning on PND 30, animals received 5 g/kg EtOH (35% w/v in H_2_O) or H_2_O via intragastric gavage (i.g.) for a total of 10 doses, using a two days on, one day off, two days on, two days off intermittent schedule over 16 days. The EtOH dosage was selected to achieve blood ethanol concentrations (BECs) consistent with those reported in adolescent humans during binge drinking episodes^26^ and our previous rodent studies^24,27^.

### AAV Packaging

All plasmids and prepackaged adeno-associated viruses (AAVs) were obtained from Addgene. AAVs containing the PHP.eB capsid were used to generate AAV-PHP.eB.gfaABC1D.Lck.GFP (Addgene plasmid #105598; RRID:Addgene_105598^28^), AAV-PHP.eB.GfaABC1D.GRAB_Ado1.0 (9.62 x 10^13^; Addgene plasmid #153288; RRID:Addgene_153288^29^), using the methods described previously by^30^. See supplemental methods section for complete packaging details. The following prepackaged AAVs were also used: AAV5.GfaABC1D.GCaMP6f (Addgene plasmid #52925; RRID: Addgene_52925^31^), AAV5.GFAP.iGluSnFr.WPRE.SV40 (Addgene plasmid #98930; RRID:Addgene_98930^32^), AAV5.GFAP.hM3D(Gq).mCherry (Addgene plasmid #50478, RRID: Addgene_50478; gift from Bryan Roth); and AAV1.hSyn.iGluSnFr.WPRE.SV40 (Addgene plasmid #98929; RRID: Addgene_98929^32^).

### Surgical Procedures

On PND 26-27, rats received 0.3 mg/kg intraperitoneal injection (i.p.) of 25% w/v mannitol^27,33,34^, buprenorphine (1.0ml/kg), and anesthetized with a cocktail of ketamine (30 mg/kg), xylazine (2.5 mg/kg), and acepromazine (0.5 mg/kg). A burr hole was drilled into both hemispheres of the skull, allowing for the bilateral injection of an AAV into the dHipp at the following coordinates, -3.12 mm AP, +/− 2.5 mm ML, -2.4 mm DV (based on the Paxinos atlas). AAVs previously discussed in ‘AAV Packaging’ were injected at 1 µL using a 28-gauge needle (cat# C313I/SPC; P1Technologies, Roanoke, VA) and syringe pump (rate: 20ml/min-330uL/sec; cat# 78-0100W; World Precision Instruments; Sarasota, FL) with a dwell time of 10 minutes. The surgical incision was closed with dissolvable, non-continuous, external sutures (cat# 662G; ETHILON® Nylon Suture; Ethicon, Raritan, NJ), and antibiotic ointment was applied to the incision. Animals were allowed to recover on a heating pad until fully awake and mobile before returning to the home cage.

#### Fiber Photometry

A subset of animals (injected with AAV1.hSyn.iGluSnFr.WPRE.SV40 or AAV-PHP.eB.GfaABC1D.GRAB_Ado1.0) were used for the fiber photometry study followed by the same surgery preparations as described above, with slight modifications allowing for fiber-optic cannula implantation. After AAV injection, a fiber (1.25 mm ferrule size; 200 μm core diameter; 0.37 NA; Neurophotometrics, San Diego, CA) was implanted into the right dHipp (-3.12 mm anterior/posterior, +/− 2.5 mm medial/lateral, -2.4 mm dorsal/ventral). Dental cement (cat# 525000 A-M Systems Sequim, WA) was used to secure the fiber-optic canula with the addition of bone anchor screws (cat# 51457; 1.59 mm O.D., 3.2 mm long; Stoelting, Wood Dale, IL).

### Immunohistochemistry (IHC)

#### Slice Preparation

All slices were prepared as previously described^27^. At PND 46 or 72, animals were deeply anesthetized with isoflurane, transcardially perfused with PBS (pH 7.4; cat# P5368, Sigma-Aldrich, St. Louis, MO) for 1 minute, followed by 7 minutes of 4% paraformaldehyde (PFA; cat# 19210, Electron Microscopy Solutions, Hatfield, PA) at a rate of 20 mL/min. Brain tissue was extracted and post-fixed in 4% PFA at 4□ overnight, followed by 30% sucrose (cat# G5516-1L, Sigma-Aldrich, St. Louis, MO) in PBS at 4 □ for 2-3 days. Brains were then placed in a plastic mold and submerged in a 2:1 solution consisting of 30% sucrose in PBS:OTC freezing compound (Electron Microscopy Sciences, Hatfield, PA), frozen and then stored at -80□. The brains were sliced on a cryostat (CM 1950, Leica Biosystems, Richmond, IL) into either 80 µm or 40 µm sections, depending on the experiment, and stored at -20 □ in a cryoprotectant (1:1 glycerol and PBS).

#### PSD-95, Ezrin, and Bassoon Immunohistochemistry

40 µm slices were rinsed three times in tris-buffered saline (TBS; pH 7.6) in 0.2% triton 100-X (TBS + 0.2% triton 100-X; TBST) and blocked at room temperature for 1 hour in 10% normal goat serum (NGS; cat# 005-000-121; Jackson Immunolabs, West Grove, PA) and TBST. Slices were incubated overnight in mouse anti-ezrin (1:500; cat# SAB4200806; RRID:AB_3714841; Sigma-Aldrich, St. Louis, MO), guinea pig anti-bassoon (1:500; cat# 141-318; RRID: AB_2927388; Synaptic Systems, Göttingen, Germany), and rabbit anti-PSD-95 (1:350; cat# 51-6900; RRID:AB_2533914; Thermo Fisher Scientific, Waltham, MA) in 10% NGS and TBST at 4 °C. Slices were washed 3 times in TBST then incubated for 2 hours at room temperature in goat anti-guinea pig 488 (1:100; Alexa Fluor cat# A11073; RRID:AB_2534117; Thermo Fisher Scientific, Waltham, MA), goat anti-mouse 594 (1:100; Alexa Fluor cat# A21125; RRID:AB_2535767; Thermo Fisher Scientific, Waltham, MA), and goat anti-rabbit Atto 647N (1:100; cat# 611-156-122; RRID:AB_10893043; Rockland Immunochemicals, Limerick, Pennsylvania) in 10% NGS and TBST. Slices were washed two times in TBST, once in TBS, and mounted on slides with MOWIOL® 4-88 Reagent (cat# 475904-100GM, Millipore, Burlington, MA) containing 1,4-diazobicyclo-2.2.2-octane (DABCO; cat# 8034560100, Sigma-Aldrich, St. Louis, MO). Slides were then left to cure in the dark for 48 hours.

#### mGluR2/3/5 and GLAST/GLT-1 Immunohistochemistry

80 µm slices were rinsed in PBST (PBS in 0.2% triton 100-X) three times. Slices underwent antigen retrieval as previously described^27,34,35^. Following antigen retrieval, slices were then blocked at room temperature for 1 hour with 5% NGS in PBST. Slices were then placed in one of the following primary antibodies combinations in 5% NGS in PBST: rabbit anti-mGluR2/mGluR3 (1:500; cat#PA1-30151; RRID:AB_1958342; Thermo Fisher Scientific, Waltham, MA) and mouse anti mGluR5 (1:500; cat# MA5-27691; RRID:AB_2735273; Thermo Fisher Scientific, Waltham, MA); guinea pig anti-GLT1 (1:1000; cat# AB1783; RRID:AB_90949; Sigma-Aldrich, St. Louis, MO) and mouse anti-GLAST (1:1000; cat# MABN794; RRID:AB_2811303; Millipore, Burlington, MA). Secondaries were incubated for 6 hours at room temperature with secondary antibodies and 5% NGS in PBST: goat anti-mouse 594 (1:200; Alexa Fluor cat# A21125; RRID:AB_2535767; Thermo Fisher Scientific, Waltham, MA) and goat anti-rabbit 647 (1:200; Alexa Fluor cat# A-21245; RRID:AB_2535813; Thermo Fisher Scientific, Waltham, MA); goat anti-mouse 594 (1:200; Alexa Fluor cat# A21125; RRID:AB_2535767; Thermo Fisher Scientific, Waltham, MA) and Alexa Fluor goat anti-guinea pig 647 (1:200; Alexa Fluor cat# A-11076; RRID:AB_141930; Thermo Fisher Scientific, Waltham, MA). Slices were washed two times in PBST, once in PBS, and mounted on slides with Vectashield containing DAPI (cat# H-1200, Vector Laboratories West Grove, PA, USA), coverslipped, and sealed with nail varnish. Slides were stored at -20 □.

### Data Acquisition and Processing

#### Astrocyte Receptors and Transporters Localization

An Olympus FV3000 laser scanning confocal microscope (Olympus, Tokyo, Japan) with an oil-immersion 40x objective was used for image acquisition (1024 × 1024 frame size, 2.3 zoom, 80um thick, optical section depth of 1 µm) of entire astrocytes of interest. A total of 6-8 randomly selected GFP^+^ individual astrocytes in CA1 dHipp (2-3 separate brain slices per animal; 5-6 brains/treatment group). The experimenter was blinded to treatment group throughout the imaging and quantification process. AutoQuant X3.1.2 software (Media Cybernetics, Rockville, MD) was used for deconvolution of the raw image files and using default parameters determined by the software algorithm and based on the confocal model, objective, and imaging specifications. Output files were directly imported to Imaris x64 version 10.2.0 (Bitplane, Santa Barbara, CA) for 3D reconstruction and quantification of astrocytes and transporter/receptor localization using methods previously described^36^. See supplemental methods for full Imaris quantification details.

#### STED PAP-synaptic Puncta Analysis and Decoupling

To obtain an overall puncta count, an Olympus FV3000 laser scanning confocal microscope (Olympus, Tokyo, Japan) with an oil-immersion 40x objective was used for image acquisition (1024 × 1024 image size, 15 µm thick, optical section depth 0.33 µm). Raw synapse count was quantified using previously published routines (https://github.com/Eroglu-Lab/Syn_Bot^37^). To assess puncta proximity, a Leica Stellaris 8 stimulated emission depletion (STED) super-resolution microscope (Cat. No. 11507584; Leica Microsystems, Germany) with an oil-immersion 93X objective was used to acquire images (1024 x 1024 image size, 5x zoom, pixels = 25nm) of the pre-, and post-synaptic terminals, and PAPs within the CA1 dHipp. Images were imported into Leica LAS X analysis software (RRID:SCR_013673; Leica Microsystems, Wetzlar, Germany). PAP-synaptic proximity was assessed using the line tool to intersect each puncta of interest creating a peak intensity plot. The following distances between the peak fluorescent intensities were measured: 1) distance between pre- and post-synaptic terminal, to determine changes in pre- and post-synaptic proximity and 2) (distance between PAP-pre-synaptic terminal + distance between PAP-post-synaptic terminal)/2, to determine the average PAP proximity to each synapse. Images were obtained from the midpoint of the stratum radiatum between the cell body pyramidal layer and the lacunosum-molecular cell body layer. Tripartite synapses were randomly selected throughout the images to remove bias (selected from evenly distributed grid sections, with each image being parsed into 9 equally sized squares). Tripartite synapses were only selected when all 3 puncta were within a distance of 1µm from each other. This distance was established through prior evaluation of tripartite synapse images obtained from EM. Quantification was conducted using 5-10 colocalization assessments/ROI from 6-10 ROIs/animal from 6 animals/trt grp.

### Physiology Recordings

#### Glutamate and Calcium Signaling Recordings

At PND 72-76, animals were deeply anesthetized with isoflurane and the whole brain was removed and placed in artificial cerebral spinal fluid (ACSF) consisting of (in mM): 125 mg NaCl, 1.6 mg KCl, 1.2 mg NaH_2_PO_4_, 2.4 mg CaCl_2_, 1.2 mg MgCl_2_, 18 mg NaHCO_3_, 11 mg glucose. 350 µm coronal sections were obtained using a Compresstome (Precisionary Instruments, Natick, MA) and slices were transferred to ACSF that was bubbling in carbogen and incubated in a water bath at 32 □ for 1 hour. Individual slices were transferred to the recording stage equipped with a Zeiss Axiocam 702 mono camera and perfused with ACSF at a constant flow rate of 2-4 ml/min maintained at 32 □. Astrocytes of interest were identified by expression of GFP (GCaMP6f or iGluSnFR). Physiological recordings were collected using methods previously described in^31^ with slight modifications. See supplemental for physiological recordings details.

#### Astrocyte Membrane Potential Recordings

Following the previously described slice physiology preparation with slight modifications, astrocytes of interest were identified by GFP^+^ expression, resting membrane potential (∼80 mV), and current-voltage properties^31^. Changes in astrocyte membrane potential were recorded following exogenous glutamate release (glutamate puff, 100µM) in the *stratum radiatum*.

### Behavioral Assays

#### Contextual Fear Conditioning

Following the 26-day forced abstinence period, animals underwent a 3-day contextual fear conditioning (CFC) paradigm previously described^2^ using Med Associates operant chambers. See supplemental for full behavioral assays. A total of six treatment-AAV groups underwent behavioral testing 30 min post-administration of clozapine-N-oxide (CNO, 2 mg/kg in 0.5% DMSO/saline, i.p.) or saline (control). Treatment groups: 1) H_2_O + AAV-PHP.eB.gfaABC1D.Lck.GFP + CNO, 2) H_2_O + AAV-PHP.eB.gfaABC1D.Lck.GFP + Saline, 3) H_2_O + AAV5.GFAP.hM3D(Gq).mCherry + Saline, 4) H_2_O + AAV5.GFAP.hM3D(Gq).mCherry + CNO, 5) EtOH + AAV5.GFAP.hM3D(Gq).mCherry + Saline, and 6) EtOH + AAV5.GFAP.hM3D(Gq).mCherry + CNO.

#### DeepLabCut

Videos were analyzed using body part tracking from DeepLabCut Software (Version 2.3.11)^38,39^ to determine freezing and darting behavior by extracting the x and y coordinates to identify movements of attempting to escape the apparatus. The number of freezing and darting responses for each animal was recorded and exported to an Excel spreadsheet. Freezing was determined by normalizing the y-axis body parts that were lowest on the apparatus and darting was determined by normalizing to the highest on the apparatus to indicate attempted escapes from the box. We averaged movement 10 sec. prior to pre-shock and 2 sec. immediately following post-shock. The pre-and post-foot shocks were defined as: pre-FS1, -FS2, -FS3 versus post-FS1, -FS2, -FS3. Code is available upon request.

### Statistical Analysis

Data were compiled using Excel 365 (Microsoft, Redmond, WA) and analyzed using GraphPad Prism Version 9.4.1 (GraphPad Software, San Diego, CA). Tripartite proximity was quantified using Student t-tests. For slice physiology analysis, custom-written software in MATLAB 2023b (The Math Works, Natick, Massachusetts) was utilized (code provided in Supplemental Fig 7) and followed by Student t-tests. For all fiber photometry data, a linear model regression analyses, mixed-effects ANOVA (timepoint x treatment) with a Tukey’s post hoc. To evaluate freezing behavior in the fear conditioning, RM two-way ANOVA, Tukey’s multiple comparisons test, n=12 rats/trt group, collapsed across 2 cohorts. The Shapiro–Wilk and Kolmogorov–Smirnov were performed to assess normality. All data were in normal distribution parameters and variance was similar between groups that were being statistically compared. Statistical significance was assessed using an alpha level of 0.05. Data are presented as violin plots with individual data points and medians.

### Data Availability

Data will be available in Distributed Archives for Neurophysiology Data Integration (DANDI) following acceptance for publication: https://dandiarchive.org.

## RESULTS

### AIE Induced Changes in PAPs-Synaptic Proximity

Our laboratory has shown that AIE produces an enduring shift toward an immature dendritic spine phenotype in the adult male rat dHipp^20^. Previous work demonstrates that dendritic spine immaturity is associated with reduced proximity of peripheral astrocytic processes (PAPs) to synapses^25^; therefore, we examined whether AIE produced similar PAP-synaptic decoupling acutely during peak withdrawal or in adulthood. Rats received AIE from postnatal day (PND) 30–45 and were euthanized either 24 h after the final ethanol dose (PND 46; peak withdrawal) or after a 26-day abstinence period (PND 72; adulthood) (Fig. 1a). Blood ethanol concentrations (190.5 ± 4.7 and 175.5 ± 4.3 mg/dL; Fig. 1b) obtained were consistent with human binge drinking levels^26^.

**Fig 1.**
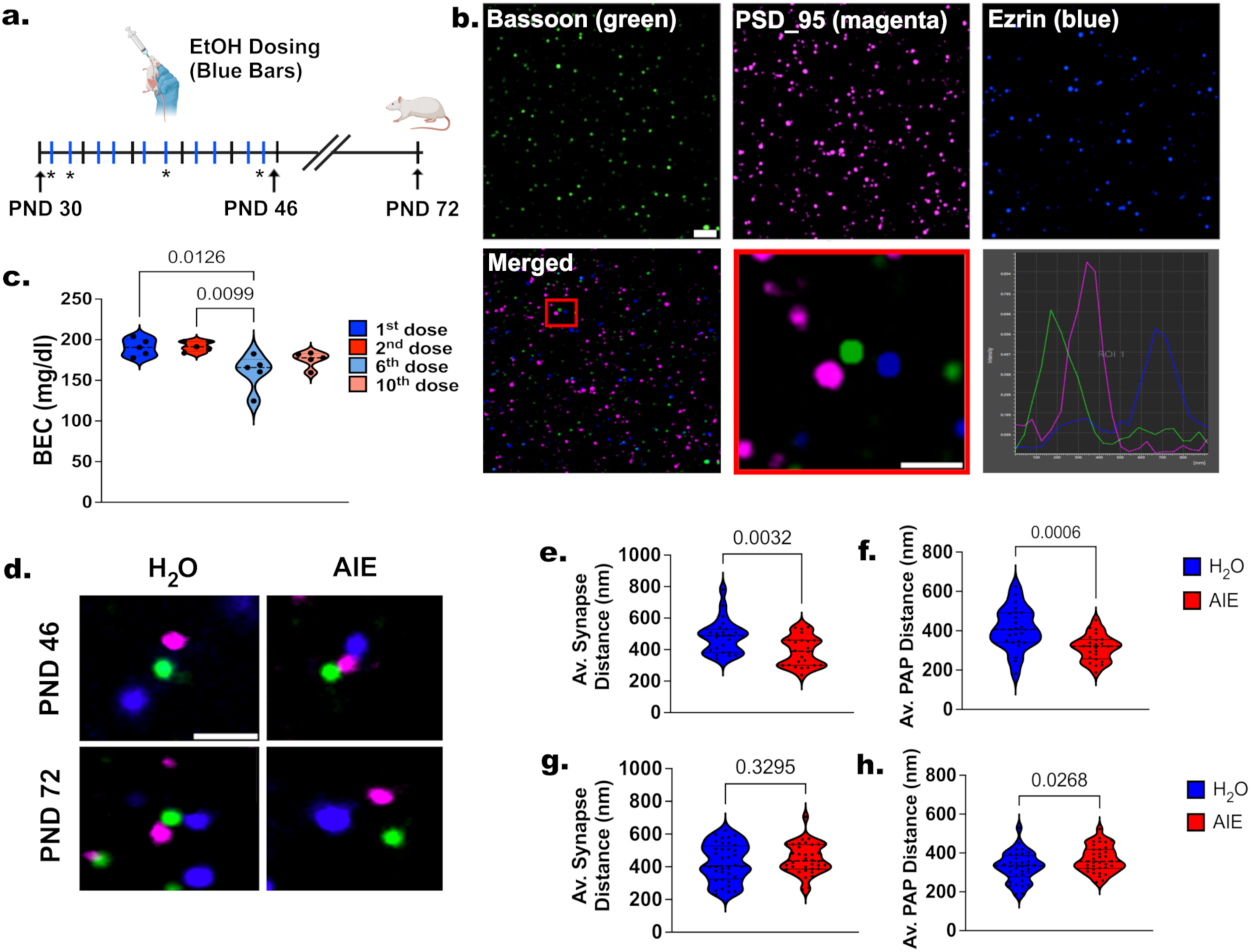
Loss of PAP synaptic proximity following AIE in adulthood. **a.** Experimental design of male adolescent rat dosing paradigm: Beginning at PND 30, animals received 5 g/kg of EtOH or H_2_O intragastrically (i.g.) on an intermittent, 2 days on, 1 day off, 2 days on, 2 days off, schedule (blue bars). Brains were collected on PND 46, 24-hr after the last dose during peak withdrawal, or after a 26-day abstinence period, on PND 72. **b.** Blood ethanol concentrations (BECs) were assessed throughout the dosing paradigm (shown as *). **c.** STED representative images of dHipp pre-synaptic bassoon (green, top left), post-synaptic PSD-95 (magenta, top center), and PAP terminal marker ezrin (blue, top right). Merged images of the tripartite synapse (bottom left and bottom center) were used to determine the distances between synaptic and PAP puncta resulting in maximum intensity plots. Distances were determined by using the absolute values between the recorded peaks (bottom right). Scale bar= 2 µm, enlarged (bottom center) scale bar= 50 µm. **d.** Representative images of tripartite synapse proximity following peak withdrawal (PND 46) and forced abstinence period (PND 72) in AIE and H_2_O exposed animals. Scale bar= 50 µm. **e, f.** Mean pre-post synaptic distance and mean PAP-synaptic distance during peak withdrawal (PND 46). **g, h.** Same measures as e,f but following the 26-day abstinence period (PND 72). Data are presented as violin plots and include the distribution of individual data points with interquartile ranges; median, minimum, and maximum measures indicated. Analysis: unpaired t-tests, n= 5 images/slide. 5-7 tripartite synapses/image. 3 slices/animal. 6 animals/treatment group.

STED microscopy of presynaptic (Bassoon), postsynaptic (PSD-95), and astrocytic (Ezrin) markers revealed that AIE decreased both pre- to post-synaptic distance (p = 0.003) and PAP-synaptic distance (p = 0.001) during peak withdrawal (Fig. 1c-f). After abstinence, pre- to post-synaptic distance returned to control levels (p = 0.330), but PAP-synaptic distance increased (p = 0.027; Fig. 1g,h). To improve the rigor of this finding, we repeated the experiment using 3D astrocyte reconstruction and quantified changed in PSD95 proximity. Using this secondary method, we demonstrated a slight reduction in astrocyte volume and confirmed the findings of the STED experiment (Supplemental Fig. 1), demonstrating a persistent loss of PAP-synaptic proximity consistent with long-term structural decoupling between astrocytes and excitatory synapses in adulthood.

### AIE-Induced Loss of PAP-synaptic Proximity Is Not Due to a Loss of Synaptic Number

We next quantified total Ezrin, Bassoon, and PSD-95 puncta and found that AIE had no effect on total Ezrin, PSD-95, or Bassoon/PSD-95 colocalized puncta (all p > 0.250; Supplemental Fig. 2 and 3b), indicating that the PAP-synaptic decoupling was not a consequence of PAP or synaptic loss.

### AIE Disrupts Astrocyte Ca^2+^ Responsivity to Neuronal Stimulation Despite Increased Glutamate Availability in Adulthood

Astrocytes remove glutamate from the synaptic cleft to prevent excitotoxicity^40^, while glutamate interactions with G-protein coupled receptors (GPCRs) located on PAPs facilitate internal Ca^2+^ (iCa^2+^) propagation, resulting in feedforward ion and gliotransmitter release^41^. Given the importance of PAP-synaptic proximity for these functions, we assessed whether PAP-synaptic structural uncoupling was sufficient to drive neuronal-to-astrocyte functional decoupling.

We targeted dorsal hippocampal (dHipp) astrocytes with an AAV5.GfaABC1D.GCaMP6f Ca² □ sensor (Fig. 2a-c). In adulthood, AIE markedly reduced astrocyte iCa²□ responses to Schaffer collateral stimulation (t-test, p = 0.040; Fig. 2d). To assess whether this impairment reflected reduced glutamate availability at the astrocyte, we expressed the membrane-bound glutamate sensor AAV5.GFAP.iGluSnFR.WPRE.SV40 on astrocytes. Baseline glutamate availability at the PAPs were unchanged (t-test, p = 0.467); however, Schaffer collateral stimulation produced enhanced glutamate-PAP interactions (t-test, p = 0.026) and slower decay (t-test, p = 0.030; Fig. 3e-g), consistent with increased extracellular glutamate availability. Additionally, the increased glutamate availability was consistent with overexpression in VGluT1, suggesting more glutamate release at the pre-synaptic terminal (t-test, n=4-6 images/rat, n=6 rats, *p*=0.0002; Supplemental Fig. 3a,b). Since astrocyte clearance of glutamate occurs via glutamate transporters (GLAST and GLT1), and glutamate-GPCR interactions at the PAPs play a vital role in the initiation of astrocyte iCa^2+^ propagation^41^, we next assessed changes in astrocytic mGluR2/3, mGluR5, GLT1, and GLAST expression. Internalized and membrane bound receptor and transporter expression remained unchanged after AIE (t-test, all p > 0.250; Fig. 3j–m, Supplemental Fig. 4). These data show that the loss of astrocyte iCa^2+^ responsivity was not due to the lack of astrocyte-specific glutamate receptor or transporter availability, suggesting that the decreased astrocyte responsivity was likely due to intrinsic astrocyte dysfunction.

**Fig 2.**
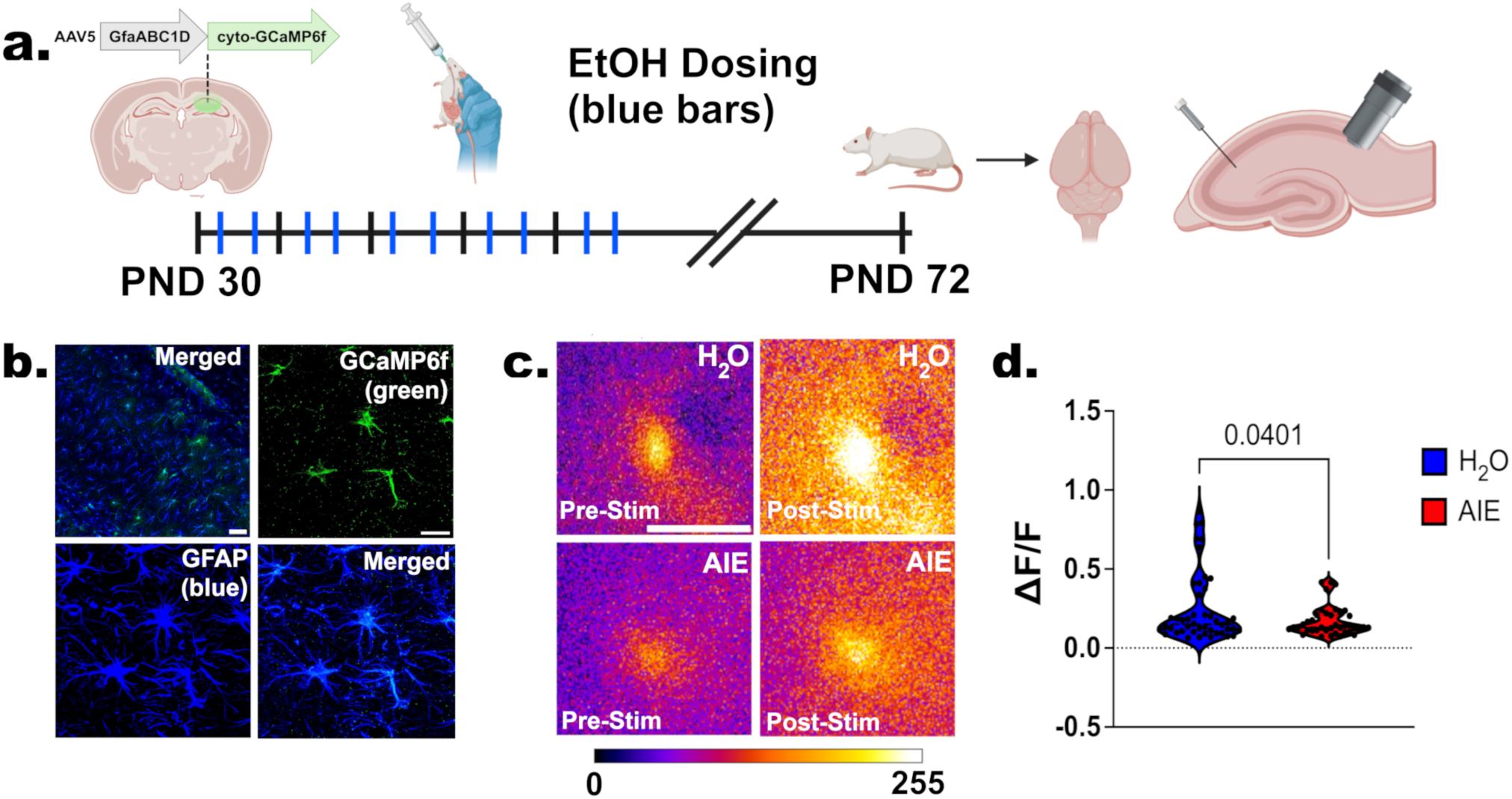
Impaired astrocyte responsivity to neuronal stimulation after AIE. **a.** Experimental design. Schematic representation of intracranial injection of the astrocyte-specific Ca^2+^ sensor (AAV5.GfaABC1D.cyto.GCaMP6f) into the dHipp on PND 26. Beginning at PND 30, animals received AIE or H_2_O intermittently through PND 45. Brains were collected on PND 72, and acute hippocampal slices were prepared for physiology. A bipolar stimulating electrode was placed in CA1 region of the *Schaffer Collaterals* to induce neuronal stimulation with simultaneous recording of astrocyte-specific Ca^2+^ signaling within the CA1 dHipp. **b.** Representative images demonstrating confirmation of GCaMP6f (green) expression and specificity for astrocytes (GFAP, blue). Scale bar = 125 µm, 2 µm. **c.** Representative images of astrocyte Ca^2+^ signaling before and after *Schaffer Collateral* stimulation following H_2_O and AIE, in adulthood. Scale bar = 10 µm. **d** Quantification of GCaMP6f signal shown as ΔF/F ratio. Data are presented as violin plots representing the distribution of individual data points. Analysis: unpaired t-tests. n = 6-10 astrocytes/animal, 6 animals/treatment group.

**Fig 3.**
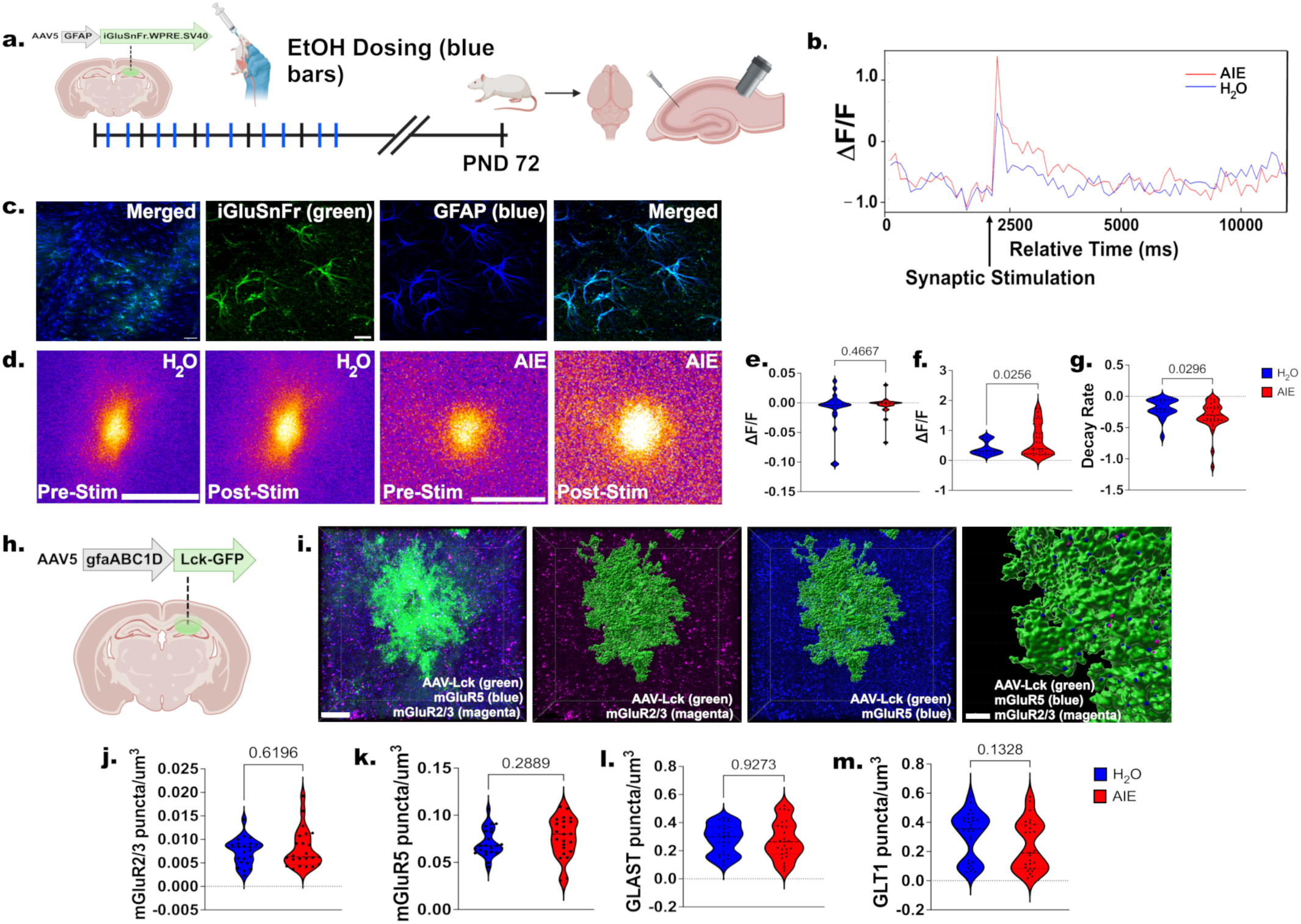
AIE-induced decrease in astrocyte responsivity to neuronal stimulation is not due to a lack of glutamate availability at the PAPs or a loss of receptor availability. **a.** Schematic representation of intracranial injection of the astrocyte-specific extracellular glutamate release sensor (AAV5.GFAP.iGluSnFr.WPRE.SV40) in the dHipp on PND 26. Beginning at PND 30, animals received AIE or H_2_O intermittently through PND 45. Brains were collected on PND 72 and prepared for acute slice physiology. A bipolar stimulating electrode was placed in CA1 region of the *Schaffer Collaterals* to induce neuronal stimulation with simultaneous recording of astrocyte-membrane specific glutamate signaling within the CA1 dHipp. **b.** Representative trace of AIE and control extracellular glutamate release following single neuronal stimulation. **c.** Representative images demonstrating confirmation of iGluSnFR (green) expression and specificity for astrocytes (GFAP, blue) within the dHipp. Scale bar =125 µm, 2 µm. **d.** Representative images of iGluSnFR in astrocytes before and after neuronal stimulation. Scale bar = 10 µm. **e.** Quantification of ΔF/F ratio prior to neuronal stimulation. **f-g**. Quantification of ΔF/F ratio and signal decay rate following neuronal stimulation. **h.** Schematic representation of intracranial injection of the astrocyte membrane-specific adeno-associated virus (AAV5.gfaABC1D.Lck.GFP) in the dHipp on PND 26. **i.** Representative images of astrocyte surface rendering (Lck-GFP, green), mGluR2/3 (magenta), and mGluR5 (blue) in the adult dHipp following AIE. Scale bar= 20 µm, 5 µm. **j-m**. Quantification of membrane-bound glutamate receptor and transporter expression. Data are presented as violin plots representing the distribution of individual data points. Analysis: unpaired t-tests, n = 6-10 astrocytes/animal, 6-8 x 80 µm image stacks/slide, 2 slices/animal, 6 animals/treatment group.

### Astrocytes Increase Astrocyte-Mediated Cation Clearance in Response to AIE

Astrocytes provide homeostatic support at the tripartite synapse through neurotransmitter uptake and the regulation of ion availability^16^. Potassium (K^+^) availability is a critical regulator of synaptic activity and, when not appropriately removed by astrocytes, can result in an increase in neuronal depolarization and hyperexcitability^42^. Based on the AIE-induced increase in glutamate availability following Schaffer Collateral stimulation, we next wanted to determine if astrocytic regulation of cation clearance, a measure predominantly driven by K^+^ flux, was upregulated. To address this question and control for glutamate release, we recorded astrocyte peak current amplitude following glutamate puff (100 µM) in the *stratum radiatum* (Fig. 4 a-c). AIE increased peak current amplitude compared to controls (t-test, n=5-9 astrocytes/rat, n=5 rats, *p*=0.025; Fig. 4d). Since the majority of K^+^ clearance occurs via Kir4.1, an inward rectifying K^+^ channel expressed predominantly at the PAP^43^ we next wanted to determine if the increase in peak current amplitude was associated with an increase in Kir4.1 expression. To do this we utilized subcellular fractionation of synaptosomes to allow us to probe for Kir 4.1 at the synapse (Supplemental Fig. 6a). Despite increased K^+^ clearance at the synapse, AIE did not alter expression of Kir 4.1 (t-test, n=5 rats/trt grp, *p*=0.301; Supplemental Fig. 6a). These results demonstrate that astrocyte-mediated clearance of K^+^ ions increases following AIE without increasing Kir 4.1 expression. This could reflect channel dysfunction or a homeostatic baseline shift in response to the chronic, excessive glutamate availability that occurs in adulthood following AIE.

**Fig 4.**
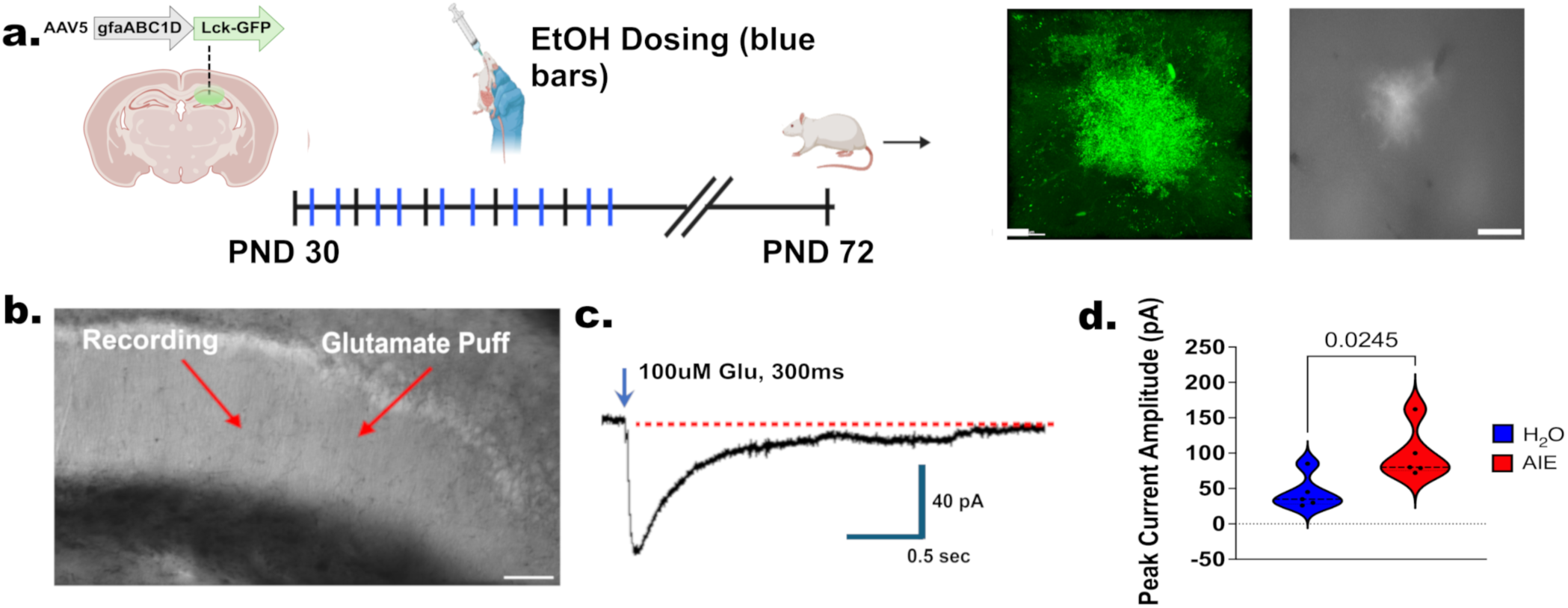
AIE results in increased astrocyte-mediated clearance of K^+^ from the tripartite synapse. **a.** Left, Schematic representation of intracranial injection of the astrocyte membrane-specific adeno-associated virus (AAV5.gfaABC1D.Lck.GFP) in the dHipp on PND 26. Beginning at PND 30, animals received AIE or H_2_O intermittently through PND 45. Brains were collected on PND 72 and prepared for acute slice physiology. Right, Representative image of astrocyte-specific AAV (Lck-GFP, green) and astrocyte excitation of GFP in the CA1 dHipp. **b.** Glutamate puff (100 µM) was used to induce neuronal stimulation and measure changes in astrocyte membrane potential in the CA1 dHipp. **c.** Representative trace of membrane potential following 100 uM glutamate puff for 300 ms. **d.** Quantification of astrocyte peak current amplitude. Analysis: unpaired t-tests,n= 6-10 astrocytes/animal. n=5-6 animals/treatment group.

### Chemogenetic Astrocyte Activation Rescues AIE-Exacerbated Fear Responding in the Contextual Fear Conditioning Paradigm

Previous work demonstrates that AIE can dysregulate dHipp-dependent behavior^2,44-47^ and that these behaviors are likely bidirectionally regulated by astrocyte iCa^2+^ signaling^48-50^. Here, we utilized a contextual fear conditioning (CFC) paradigm, an experimental model of fear learning in which, animals learn to associate a neutral context with an aversive stimulus. Following acquisition and consolidation, animals will display fear responses to the context that predicts danger which will be distinguished over time if no aversive stimulus is presented. Changes in acquisition can provide indications of heightened fear responsivity and an inability to learn fear, both of which can result in compromised decision making and mental health outcomes in humans. To determine whether restoring astrocyte iCa^2+^ activity is sufficient to recover AIE-induced heightened fear responding during the acquisition phase of the CFC task, we expressed astrocyte-specific designer receptor activated by designer drugs (DREADDs, AAV5.GFAP.hM3D(Gq).mCherry) in the dHipp and administered clozapine-N-oxide (CNO, 2 mg/kg, i.p.) 30 min. prior to the CFC day 1 (Fig. 5a, b). Freezing and darting behavior were evaluated throughout all 3 sessions using DeepLabCut (Fig. 5c). Additional behaviors were assessed to ensure no off-target effects of CNO (Supplemental Fig. 7).

**Fig 5.**
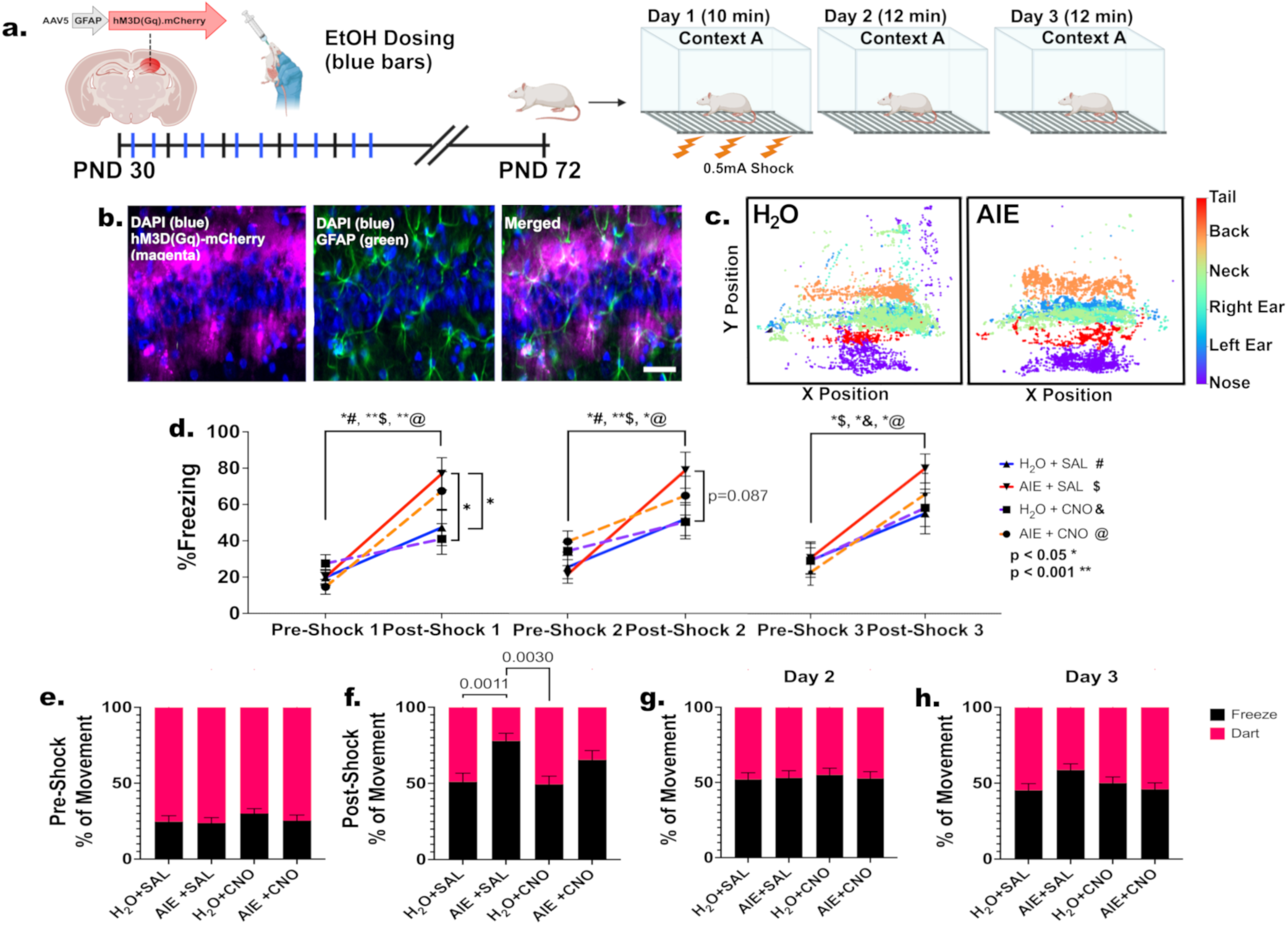
Chemogenetic recovery of astrocyte activity is sufficient to partially rescue dHipp-induced behavioral deficits in fear learning acquisition following AIE. **a.** Schematic representation of intracranial injection of the astrocyte membrane-specific adeno-associated virus (AAV5.GFAP.hM3D(Gq).mCherry) in the dHipp on PND 26. Beginning at PND 30, animals received AIE or H_2_O intermittently through PND 45. Following 26 day forced abstinence period, animals received saline or clozapine-N-oxide (CNO, i.p. 2 mg/kg) to activate astrocytes. After 30 minutes, animals were assessed in a contextual fear conditioning paradigm. Animals were placed in context A and subjected to 3-foot shocks across a 10-minute test period on day 1. On day 2 and 3, animals were placed in context A for 12 min with no foot shocks. **b.** Representative images of the dHipp, DAPI (blue), hM3DGq (magenta), GFAP (green), and colocalization of GFAP+hM3DGq confirming astrocyte specificity of AAV. **c.** Representative images of DeepLabCut output of labeled body parts on the x-y axis of the context box. **d.** Quantification of freezing behavior in the CFC task (H_2_O+SAL (#) vs. AIE+SAL ($); *p = 0.029-0.085;* H_2_O+CNO (&) vs. AIE+CNO (@); *p = 0.062-0.574*). **e-f**. Freeze/dart behavior quantification from sessions collapsed across foot shocks from Day 1 of the CFC. **g-h.** Freeze/dart behavior quantification from Days 2 and 3 of the CFC. Analysis: Repeated measures Two-way ANOVA, n = 12 animals/treatment group across 2 separate cohorts. (*p < 0.05* p < 0.001**)*

#### Acquisition Phase (day 1)

Pre- and post-foot shocks were defined as: pre-FS1, -FS2, -FS3 versus post-FS1, -FS2, -FS3. During pre-FS1, there were no baseline differences in freezing behavior when assessed across treatment groups (RM two-way ANOVA, Tukey’s multiple comparisons test, n=12 rats/trt group, collapsed across 2 cohorts, average *p* > 0.05; Fig. 5d). Post-FS1, AIE animals displayed significantly higher freezing behavior compared to controls (H_2_O+SAL vs. AIE+SAL; RM two-way ANOVA, Tukey’s multiple comparisons test, *p*=0.029; Fig. 5d); this AIE effect was subtly attenuated by chemogenetic astrocyte activation (H_2_O+CNO vs. AIE+CNO; *p*=0.062). Despite AIE-induced heightened freezing in response to FS1, when freezing was re-assessed immediately prior to FS2 (i.e., pre-FS2), all animals showed recovery to similar baseline freezing levels (all *p’s* = 0.377-0.967). During post-FS2, AIE animals showed a trend towards higher freezing behavior when compared to controls (H_2_O+SAL vs. AIE+SAL; RM two-way ANOVA, Tukey’s multiple comparisons test, *p*=0.085); however, there were no differences between all other treatment groups (all *p’s* = 0.599-0.999). Consistent with the patterns observed at pre-FS1 and pre-FS2, all animals returned to comparable baseline freezing levels by pre-FS3 (all *p’s* = 0.934-1.00). By post-FS3, there were no differences between groups (all *p’s* = 0.246- 0.996). These data indicate that AIE animals exhibit initial heightened freezing responses to foot shock, which subsequently normalized because of the other groups’ more gradual increases in freezing across foot shocks. To assess generalized changes in freezing behavior across the session, the data were collapsed across all three-foot shocks and changes in % movement, categorized as freezing (<50% movement) and darting (> 50% movement), were assessed between treatment groups (Fig. 5e-h). Once again, during pre-FS1-3, there were no differences between treatment groups (pre-S1-3, all *p’s* =0.484-0.999). During post-FS1-3, AIE animals showed increased freezing behavior and a reduction in darting behavior when compared to controls (RM two-way ANOVA, Tukey’s multiple comparisons test, H_2_O+SAL vs. AIE+SAL *p*=0.0011; H_2_O+CNO vs. AIE+SAL *p*=0.003). AIE animals that received chemogenetic activation showed a modest rescue of freezing behavior as demonstrated by a loss of significance when compared to the H_2_O+SAL group (H_2_O+SAL vs. AIE+CNO; p=0.214) despite not reaching significance when compared to AIE+SAL (AIE+SAL vs. AIE+CNO; p=0.4014).

#### Retention and Extinction (day 2) and Extinction (day 3)

Despite AIE-induced increased freezing behavior during fear acquisition, there were no differences in retention or extinction during day 2 or 3 (RM two-way ANOVA, Tukey’s multiple comparisons test, all *p’s*= 0.124-1.000). These data show that chemogenetic astrocyte activation partially attenuates AIE-induced heightened fear responding during CFC acquisition. The lack of AIE effect on fear recall and fear extinction further suggests that astrocytes may play an important role in punishment responsivity rather than CFC task learning per se.

### Chemogenetic Astrocyte Activation Rescues AIE-Induced Decoupling of Adenosine and Freezing Behavior During the Contextual Fear Conditioning Paradigm

It has been well established that chemogenetic astrocyte activation using hM3DGq+CNO involves the upregulation of astrocyte iCa^2+^, and that the propagation of this signaling can lead to an array of downstream events^51^ (see^52^ for review), including the release of the gliotransmitter adenosine: an important modulator of neuronal activity^53^ and fear consolidation through Hipp mechanisms^50^. Based on these previous findings, we wanted to determine if the AIE-induced heightened fear responding was associated with a disruption of downstream adenosine signaling and, if so, whether chemogenetic rescue of astrocyte activity was sufficient to recover functional coupling between freezing behavior and adenosine signaling. To address these questions, we co-injected an astrocyte-specific adenosine sensor (AAV-PHP.eB.GfaABC1D.GRAB_Ado1) and astrocyte-specific hM3DGq (AAV5.GFAP.hM3D(Gq).mCherry) into the dHipp and inserted an optical fiber into the same region of the dHipp and administered clozapine-N-oxide (CNO, 2 mg/kg, i.p.) 30 min. prior to the CFC day 1 (Fig. 6a, b). During pre-FS1-3, animals that received H_2_O or AIE followed by astrocyte activation (+CNO) showed significantly higher adenosine availability when compared to non-activated (+SAL) groups (RM two-way ANOVA, Tukey’s multiple comparisons test, n=6 rats/trt group, H_2_O+SAL vs. H_2_O+CNO *p*=0.049; AIE+SAL vs. AIE+CNO *p*=0.010, Fig. 6c). At post-FS1-3, only AIE+CNO showed persistently higher adenosine availability when compared to AIE+SAL and H_2_O+CNO groups (RM two-way ANOVA, Tukey’s multiple comparisons test, AIE+SAL vs. AIE+CNO *p*=0.010; H_2_O+CNO vs. AIE+CNO *p*=0.004). To determine the relationship between adenosine availability and freezing behavior we conducted linear regression during fear acquisition (Linear regression analysis, reference= H_2_O+SAL, model fit=*R^2^*=0.270, Adj. R^2^=0.205, *F* (7,73) =3.940, *p=*0.001). When animals received H_2_O+SAL there was a positive correlation between adenosine availability and freezing behavior; however, this association was decoupled following AIE (AIE+SAL, β= - 0.013, *SE*= 0.004, *p* < 0.001, *R^2^*=0.268). When AIE animals received astrocyte activation (AIE+CNO), the AIE-induced negative association between adenosine availability and freezing behavior was completely attenuated and indistinguishable from controls (H_2_O+CNO, β= 0.530, *SE* = 0.206, *p* = 0.012, *R^2^*=0.023; AIE+CNO β = 0.346, *SE* = 0.220, *p* = 0.120, *R^2^*=0.106; Fig. 6e). These data show that chemogenetic dHipp astrocyte activation was sufficient to reverse AIE-induced decoupling of adenosine signaling and freezing behavior during fear acquisition. Interestingly, when membrane and internalized adenosine receptor and transporter expression were quantified following AIE, there was a significant increase in membrane-bound adenosine receptor A2a and adenosine transporter ENT2 (t-test, n=5-9 astrocytes/rat, n=5 rats, ENT2; *p*=0.046, A2a; *p*=0.002), but no changes in the expression of the inhibitory A1 receptor or ENT1 (t-test) when compared to controls (Supplemental Fig. 8). These data demonstrate a complex interplay of adenosine and receptor availability following AIE and astrocyte chemogenetic activation that are sufficient to partially recover appropriate fear responsivity.

**Fig 6.**
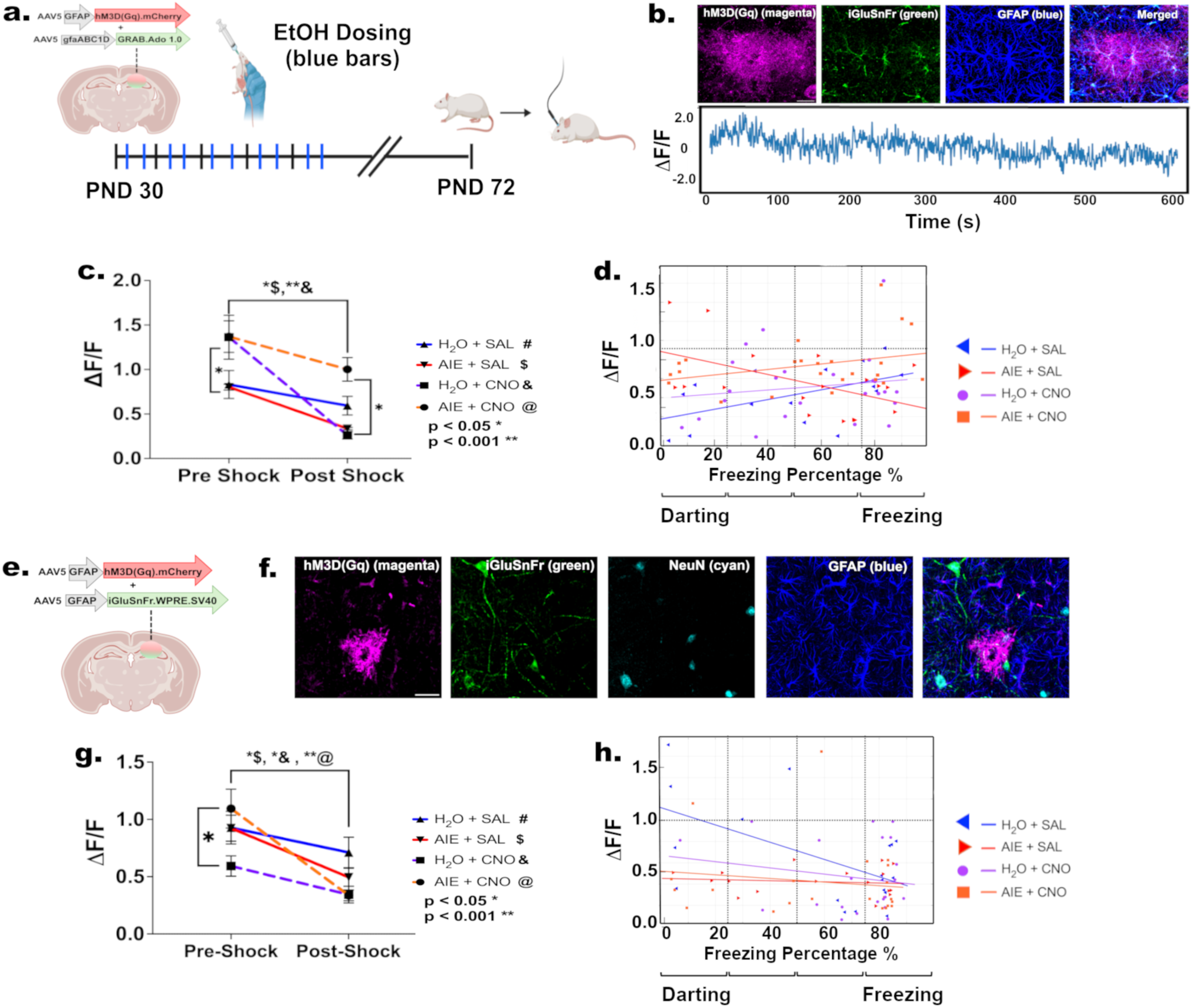
Astrocyte activation partially rescues adenosine-freezing functional coupling following AIE. **a.** Schematic representation of intracranial injection of the astrocyte membrane-specific adeno-associated virus (AAV5.GFAP.hM3D(Gq).mCherry + AAV-PHP.eB.GfaABC1D.GRAB_Ado1) and a fiber implant in the dHipp on PND 26. Beginning at PND 30, animals received AIE or H_2_O intermittently through PND 45. Following 26 day forced abstinence period, animals received saline or clozapine-N-oxide (CNO, 2 mg/kg, i.p.) to activate astrocytes. After 30 minutes, animals were then assessed for cognitive performance in a contextual fear conditioning paradigm along with real-time glutamate or adenosine availability measurements. **b.** Representative images of the dHipp, hM3DGq (magenta), GRAB.Ado1.0 (green), GFAP (blue), and colocalization of GRAB.Ado1.0+hM3DGq confirming astrocyte specificity of AAV and fiber photometry representative trace across day 1 of fear acquisition. **c.** Quantification of the adenosine ΔF/F ratio before and after foot shock: Pre-FS: (# vs. & ;*p=*0.0499; $ vs. @; *p=*0.0103) Post FS1-3: ($ vs. @; *p=*0.0103; & vs. @; *p=*0.0041). **d.** Scatterplot and trendlines comparing adenosine ΔF/F with freezing behavior (H_2_O+SAL, β= 0.007, *SE*= 0.003, *p* = 0.015, *R^2^*=0.486; AIE+SAL, β=-0.013, *SE*= 0.004, *p* < 0.001, *R^2^*=0.268; H_2_O+CNO, β= -0.005, *SE* = 0.004, *p* = 0.012*, R*^2^=-0.029; AIE+CNO, β = -0.004, *SE* = 0.003, *p* = 0.120, *R^2^*=-0.044). **e.** Schematic representation of intracranial injection of the astrocyte membrane-specific adeno-associated virus (AAV5.GFAP.hM3D(Gq).mCherry + AAV5.hSyn.iGluSnFr.WPRE.SV40) **f.** Representative images of the dHipp, hM3DGq (magenta), iGluSnFr (green), GFAP (blue), and localization of iGluSnFr and hM3DGq within a dHipp astrocyte and neuron. **g.** Quantification of glutamate availability (ΔF/F ratio) before and after foot shock (&; *p=0.0166*). **h.** Scatterplot and trendlines comparing glutamate availability ΔF/F and freezing behavior (H_2_O+SAL, β= -0.008, *SE* = 0.0024, *p* = 0.001, *R^2^*=0.319; H_2_O+CNO, β= 0.005, *SE* = 0.004, *p* = 0.229, *R^2^*=0.034; AIE+SAL, β =0.008, *SE* = 0.004, *p* = 0.047, *R^2^*=0.015; AIE+CNO, β =0.006, *SE* = 0.225, *p* = 0.066, *R^2^*=0.020). Analysis: Repeated measures Two-way ANOVA and Linear Regression Analysis, n = 6 animals/treatment group. (*p < 0.05* p < 0.001**)*

### Chemogenetic Astrocyte Activation and AIE Independently Decouple Glutamate Signaling and Freezing Response with no Additive or Synergistic Effects

Our findings indicate that chemogenetic activation of astrocytes is sufficient to restore downstream adenosine availability and re-establish its association with freezing behavior during fear acquisition. Building on this demonstration of gliotransmitter functional recovery, we next investigated whether chemogenetic restoration of astrocyte activity could likewise rescue synaptic glutamate signaling disrupted by AIE. To address this question, we co-injected a neuron-specific glutamate (AAV1.hSyn.iGluSnFr.WPRE.SV40) sensor and an astrocyte-specific hM3DGq (AAV5.GFAP.hM3D(Gq).mCherry) into the dHipp along with optical fiber implantation to obtain fiber photometry measurements during CFC (Fig. 6f, g). Animals received CNO/saline injection 30 min. prior to the CFC Day 1.

#### Acquisition Phase (day 1)

Glutamate measurements were collapsed across pre-FS1-3 and post-FS1-3 (Fig. 6h). During pre-FS1-3 (baseline), animals that received AIE followed by astrocyte activation (AIE+CNO) showed significantly higher glutamate availability when compared to the control group (H_2_O+CNO) (RM two-way ANOVA, Tukey’s multiple comparisons test, n=6 rats/trt group, H_2_O+CNO vs. AIE+CNO *p*=0.017). Post FS1-3, glutamate synaptic availability declined when compared to pre FS1-3; however, there were no differences between treatment groups (all *p’s* = 0.096-1.0000). To assess general glutamate availability, data were collapsed across the session. Unlike the adenosine experiment, there were differences in glutamate availability across treatment groups (one-way ANOVA, Tukey’s multiple comparisons test; all *p’s*=0.415-0.999; Fig. 6i). Lastly, we conducted a linear regression to assess the association between glutamate availability and freezing behavior throughout the session (Linear regression analysis, reference= H_2_O+SAL, model fit=*R^2^*=0.205, Adj. R^2^=0.125, *F* (7,70) =2.58, *p=*0.020). Within the control group (H_2_O+SAL), there was a robust negative correlation between glutamate availability and freezing behavior (β= -0.008, *SE* = 0.002, *p* = 0.001, *R^2^*=0.319; Fig 6j). However, this association was lost following astrocyte activation and following AIE administration (H_2_O+CNO, β= 0.005, *SE* = 0.004, *p* = 0.229, *R^2^*=0.034; AIE+SAL, β =0.008, *SE* = 0.004, *p* = 0.047 *R^2^*=0.015; Fig. 6j). Interestingly, when AIE and astrocyte activation were combined (AIE+CNO), there were no additive or synergistic effects (AIE+CNO, β=0.006, SE= 0.003, *p*=0.066, *R^2^*=0.021). Thus, both AIE and astrocyte activation independently uncouple synaptic glutamate signaling from behavioral output, in contrast to the restored adenosine–behavior coupling observed under astrocyte activation. These results demonstrate that glutamate availability has an inverse relationship with freezing behavior during fear acquisition and that this interaction is disrupted following AIE. Moreover, chemogenetic astrocyte activation is not sufficient to restore the negative association between synaptic glutamate availability and freezing behavior during fear acquisition.

## Discussion

The goal of this study was to define how AIE exposure influences astrocyte-synaptic communication within the dHipp and is associated with enduring dysregulation in fear learning and extinction into adulthood. We demonstrate that AIE disrupts the structural and functional integrity of the tripartite synapse by driving a persistent loss of PAP-synaptic coupling. This decoupling coincides with reduced astrocyte iCa²□ responsivity to neuronal stimulation, despite increased glutamate availability at PAPs, and manifests as exaggerated fear responding during acquisition. Remarkably, selective chemogenetic activation of dHipp astrocytes was sufficient to partially attenuate AIE-induced heightened fear responses and fully restore the positive association between adenosine signaling and freezing behavior during fear acquisition. These findings reveal that AIE drives long-lasting astrocyte-specific pathology that persists into adulthood after abstinence, and that the associated behavioral dysfunction and downstream impaired gliotransmitter release can be overcome through the targeted modulation of astrocyte activity.

### Astrocyte structural plasticity and tripartite remodeling

Astrocytic processes are highly dynamic regulators of the synaptic microenvironment. Their intimate contact with excitatory synapses is essential for maintaining ionic and neurotransmitter homeostasis and for fine-tuning synaptic efficacy through iCa²□-dependent feedforward signaling. We show that AIE induces biphasic remodeling of PAP-synaptic proximity defined by an early transient increase in proximity followed by a protracted loss of proximity during abstinence. This pattern of acute increased coupling followed by chronic decoupling in adulthood likely reflects an initial compensatory attempt to regulate glutamate availability and stabilize hyperexcitable circuits in response to repeated cycles of EtOH withdrawal, this would be consistent with previous work showing increased cortical astrocyte Ca^2+^ activity during withdrawal^54,55^. Chronic decoupling observed during adulthood, could be driven by a number of factors; 1) a failure of synapses to mature without major synaptic loss, resulting in heightened synaptic plasticity (both of which we have previously observed^20^), suggesting that PAP withdrawal is a consequence of tripartite destabilization rather than synapse elimination. 2) intentional PAP retraction that would limit astrocytic damage due to chronic excessive glutamate availability or, 3) possible microglial phagocytosis of PAPs, as has previously been observed in rat models of intermittent access to cocaine^56^. Importantly, the loss of PAP-synaptic coupling mirrors glial remodeling that has been observed across multiple disease domains, including substance use^57^, stress^58^, Alzheimer’s disease^59^, and epilepsy^25^. This convergence supports the premise that repeated insult drives persistent astrocytic maladaptive structural plasticity that outlasts the initial exposure and emphasizes an ongoing need to further understand how astrocyte maladaptive remodeling contributes to synaptic dysregulation across a variety of disease states.

### Astrocyte hypoactivity and ion-channel adaptation

The protracted loss of PAP-synaptic structural proximity suggests a functional decoupling between neurons and astrocytes. While we did observe a loss of astrocyte iCa^2+^ signaling following AIE, this was despite increased glutamate vesicle transporter expression within the pre-synaptic terminals and glutamate interactions at the PAP, supporting prior reports in AIE drives maladaptive neuronal hyperexcitability^20,60,61^. In this AIE model, astrocytes appear to counteract this hyperactivity through enhanced K□ clearance, reflected by increased inward-rectifying currents^62-64^. Interestingly, we did not observe increased Kir 4.1 expression, suggesting that the enhanced K^+^ clearance is likely a reflection of increased functional plasticity at the level of channel function, indicative of changes in gating kinetics, subcellular localization, or heteromeric assembly^65-67^. While this may be an indication of a chronic compensatory increase in basal K^+^ clearance, this does not appear to be reflected in all aspects of astrocyte function following AIE. The absence of changes in glutamate receptor or transporter expression points to a fundamental shift in strategy, away from receptor trafficking and toward ion-channel regulation, as a means of modulating local excitability. To our knowledge, this represents the first evidence that AIE produces long-term functional remodeling of astrocyte physiological responses to neuronal input.

### Astrocyte Ca²□ hypoactivity as a mechanistic target

Despite increased glutamate availability, astrocytes failed to mount appropriate iCa²□ responses following AIE. This pattern mirrors those seen in aging and neurodegeneration^68,69^ and suggests that AIE induces a premature or maladaptive “senescent-like” state in astrocytes^70^. Such iCa²□ hypoactivity may fundamentally alter astrocyte-neuron bidirectional coordination within the dHipp, impairing the encoding of contextual and fear-related memories^71,72^. Indeed, we found that AIE animals exhibited heightened fear responding during acquisition, replicating earlier work^73^. Furthermore, restoring astrocyte iCa²□ dynamics via chemogenetic activation was sufficient to partially attenuate heightened fear responding. These behavioral deficits closely resemble post-traumatic stress disorder (PTSD) where the patients exhibit exacerbated fear responses to trauma ‘triggers’ and fail to recognize the removal of the threat^74^. These results provide the first direct evidence that dHipp astrocyte activity contributes to fear responding during the contextual fear conditioning task, and that disruption of astrocyte activity acts as a mediator of AIE-induced behavioral impairment.

While the work conducted here focuses on the role of excitatory synapses, it is well established that glutamatergic and GABAergic signaling work together with astrocytes to maintain excitation/inhibition (E/I) balance^75^. Previous work has shown that AIE produces long-lasting disruption to neuronal glutamatergic signaling, GABA receptor expression, and inhibitory signaling into adulthood^20,60,76,77^. However, astrocytes also express functional NMDA and GABA receptors that contribute to the regulation of iCa²□ dynamics^75,78,79^. Further work is needed to determine how alcohol-induced GABAergic dysfunction may contribute to the reduced astrocytic iCa²□ signaling observed here^80-82^. Based on these findings, it is possible that AIE may impair astrocyte activity through broader dysregulation of both excitatory and inhibitory neurotransmitter systems, rather than through glutamatergic mechanisms alone.

### Adenosine signaling as a downstream effector of behavioral dysregulation

Fiber photometry revealed tight coupling between adenosine release and freezing behavior that was abolished by AIE and that was subtly reinstated through astrocyte chemogenetic activation. It is interesting that AIE alone did not alter baseline (pre-shock) adenosine; however, astrocyte activation in AIE animals significantly increased adenosine levels, combined with AIE-induced increases in A2a receptor expression, would suggest that enhanced adenosine availability and A2a receptor availability is necessary to recover appropriate freezing responses. This is consistent with previous work supporting a role for A_2A_ receptors in fine-tuning synapses. For example, upregulation of A2a has been associated with increased neuronal excitability and neurodegeneration (reviewed in^83^) as well as behavioral impairments specifically associated with astrocyte Ca^2+^ hypoactivity^84,85^. However, the physiological characteristics of adenosine receptor biology are quite complex. Presynaptic A1-A2a heterodimers have been shown to inhibit A1-mediated reductions in glutamate release within striatal areas^86^ and provides a mechanism to explain the paradoxical observation of high-level adenosine availability following astrocyte activation, increased glutamate release, and the changes in long-term potentiation (LTP) we have previously observed following AIE^20,87^. The unusual interplay of adenosine receptors and subunit dimerization could further explain why glutamate availability remains high following astrocyte activation. Continued excessive glutamate availability, despite astrocyte activation, may explain why there is only a partial restoration of fear responsivity. Alternatively, the associated behavioral changes within the CFC task could reflect the engagement of a novel astrocyte-driven pathway and not the rescue of regulatory pathways that typically oversee fear responsivity during foot shock acquisition. While these data establish a functional link between astrocyte-derived adenosine and freezing behavior during acquisition, the precise mechanistic contributions of astrocyte signaling during fear conditioning remains to be defined.

Together, these findings position adenosine signaling as a downstream effector of astrocyte dysfunction, linking molecular changes at the tripartite synapse to circuit-level hyperexcitability and behavioral dysregulation of fear responsivity. These findings suggest that impaired astrocyte iCa²□ signaling disrupts purinergic feedforward signaling critical for adaptive fear responding during acquisition. The current work provides important insight into the role of astrocytes in the regulation of dHipp-dependent behavioral outcomes while acknowledging that robust chemogenetic Ca^2+^ activation does not entirely recapitulate physiological events.

### Broader implications and translational potential

Although chemogenetic activation restored astrocytic activity and partially attenuated AIE-induced behavioral dysregulation, it did not fully recover glutamate homeostasis. Dissecting circuit-specific astrocyte-neuron ensembles, particularly those coordinating dHipp-prefrontal cortex communication, will be essential for identifying when and how astrocytes constrain or facilitate adaptive behavior^88-90^. Emerging tools, such as real-time fiber photometry-based functional recordings of astrocyte and neuronal activity combined with machine-learning-based behavioral mapping, now offer an unprecedented opportunity to decode these interactions *in vivo*. However, a great deal of work is still required to further our understanding of how the multitude of neurotransmitters and gliotransmitters converge to fine-tune synaptic and circuit activity and regulate subsequent behavioral outcomes across disease states.

Collectively, our results establish astrocyte structural and functional plasticity as key determinants of post-AIE circuit remodeling. They extend the concept of “glial pathology” beyond traditional neurodegeneration to include adolescent alcohol exposure as a driver of long-lasting tripartite synapse dysfunction. By revealing that these astrocytic deficits are reversible through targeted activation, this work opens a new therapeutic avenue for normalizing network excitability and improving mental health outcomes after early-life alcohol misuse.

## Conclusion

This study identifies astrocyte-specific structural and functional remodeling as a persistent cellular signature of adolescent alcohol exposure. AIE drives enduring decoupling of the tripartite synapse, disrupts astrocytic iCa²□ and adenosine signaling, and heightens fear-related behavior; yet these effects remain amenable to astrocyte-targeted intervention. Recognizing astrocytes as active, plastic regulators of neural circuits reframes them from passive supporters to key therapeutic entry points for treating alcohol- and stress-related neuropsychiatric disease.

## Supporting information

Supplemental Results

## Acknowledgments

Special thanks to Katherine Reissner (University of North Carolina, Chapel Hill) for the generous gift of the packaged AAV2/5.gfaABC1D.Lck.GFP (Addgene plasmid #105598; RRID:Addgene_105598^28^), and Isabella Parsons for her aid in immunohistochemistry staining and Imaris analysis.

