## Supplemental Results for "ADOLESCENT ALCOHOL EXPOSURE DISRUPTS ASTROCYTE-SYNAPTIC STRUCTURAL AND FUNCTIONAL COUPLING IN THE MALE DORSAL HIPPOCAMPUS"

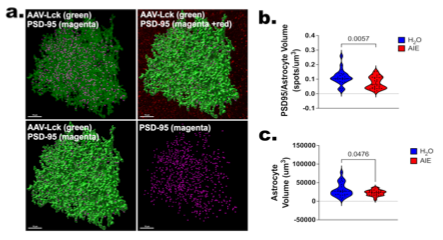
**Supplemental Fig 1. AIE-induced loss of PAP-synaptic proximity in adulthood replicated in 3D astrocyte reconstruction method. a.** Representative images of control astrocyte surface rendering (Lck-GFP, green) and PSD-95(magenta) in the top left image, top right: astrocyte surface rendering (Lck-GFP, green) with PSD-95 (magenta) within 0.5 µm of the astrocyte and PSD-97 not colocalized (red), bottom left image: PSD-95 (magenta) within 0.5 µm of the astrocyte surface rendering (Lck-GFP, green): bottom right image is isolated PSD-95 (magenta) that is colocalized with the astrocyte of interest. Scale bar= 10 µm. **b.** Quantification of puncta/astrocyte volume (puncta/µm^3^). **c.** Quantification of astrocyte volume (µm^3^). (Data are presented as violin plots and include the distribution of individual data points. Analysis: unpaired t-test, n = 6-10 astrocytes/animal, 6-8 x 80 µm image stacks/slide, 2 slices/animal, 4 animals/treatment group.

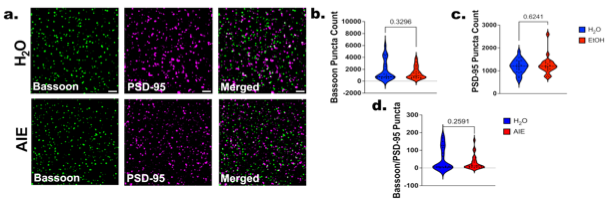
**Supplemental Fig 2. AIE-induced loss of PAP-synaptic proximity in adulthood is not due to a loss of excitatory synapses. a.** Representative images of dHipp pre-synaptic bassoon (green), post-synaptic PSD95 (magenta), and colocalized bassoon-PSD95 (white; white arrows) under H_2_O and AIE conditions, during abstinence (PND 72). Scale bars = 2 µm. **b-d.** Quantification of pre-synaptic bassoon, post-synaptic PSD95, and co-localization. Data are presented as violin plots and include the distribution of individual data points. Analysis: unpaired t-tests. All data are presented as average puncta count/image. n=3 x 15 µm image stacks/slide, 2 slices/animal, 6 animals/treatment group.

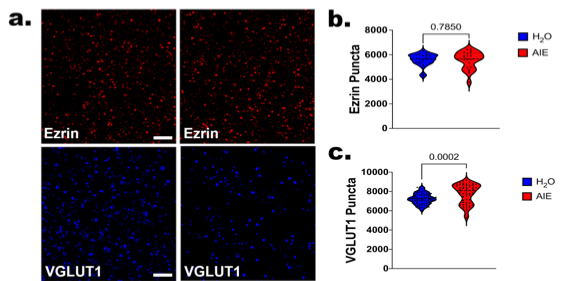
**Supplemental Fig 3. AIE-induced loss of PAP-synaptic proximity in adulthood is not due to a loss of PAPs, but AIE does increase VGLUT1 expression. a.** Representative images of dHipp PAP marker Ezrin (red) and VGLUT1 (blue) pre-synaptic excitatory vesicular marker under H_2_O and AIE conditions, following abstinence (PND 72). Scale bars = 2 µm. **b-c.** Quantification of VGLUT1 and Ezrin puncta. Data are presented as violin plots and include the distribution of individual data points. Analysis: unpaired t-tests. All data are presented as average puncta count/image. n=3 x 15 µm image stacks/slide, 2 slices/animal, 6 animals/treatment group.

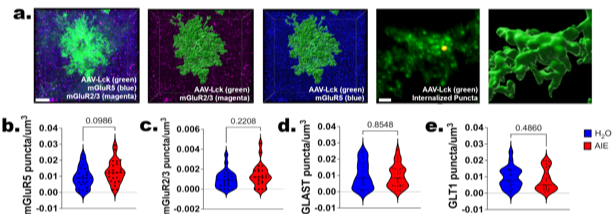
**Supplemental Fig 4. Impaired astrocyte responsivity to neuronal stimulation is not due to the internalization of glutamatergic receptors. a.** Representative images of astrocyte surface rendering (Lck-GFP, green) and internalized mGluR2/3 (magenta), and mGluR5 (cyan) in the dHipp following AIE in adulthood. Scale bar= 5, 20 µm. **b-e**. Quantification of glutamate transporter and receptor expression after AIE. Data are presented as violin plots representing the distribution of individual data points. Analysis: unpaired t-tests, n = 6-10 astrocytes/animal, 6-8 x 80 µm image stacks/slide, 2 slices/animal, 6 animals/treatment group.

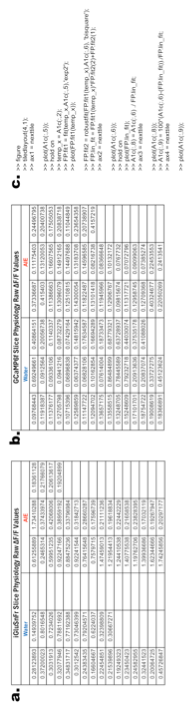
**Supplemental Fig 5. Astrocyte-specific GCaMP6f and iGluSnFr Summary. a, b.** Table representing the ΔF/F values (z-score) obtained from the astrocyte-specific GCaMP6f and iGluSnFr in response to neuronal stimulation in Schaffer Collaterals of CA1 dHipp. **c.** Coding performed on the raw fluorescence that was normalized to account for photobleaching by calculating the ΔF/F (z-score) and to determine changes in fluorescent intensity.

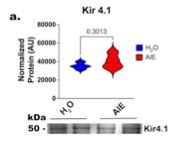

**Supplemental Fig 6. AIE did not alter expression of Kir 4.1 in the hippocampus. a.** No AIE-induced changes in Kir 4.1 expression. Representative images of Western blot probing for Kir4.1 in synaptosome isolated lysates prepared from Hipp tissue. No AIE-induced changes in Kir 4.1 expression. Analysis: unpaired t-tests, n=5 animals/treatment group.

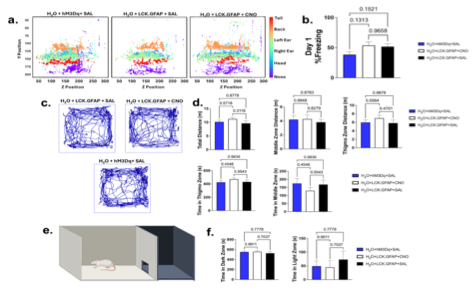
**Supplemental Fig 7. CNO produces no off-target behavioral effects in the contextual fear acquisition, open field, or light-dark tasks.** H_2_O+hM3Dq+SAL, H_2_O+LCK.GFAP+CNO and H_2_O+LCK.GFAP+SAL groups were assessed to ensure that there were no viral or CNO off-target effects. **a.** Representative images of DeepLabCut output of labeled body parts on the x-y axis in the operant chamber during CFC. **b.** Quantification of freezing behavior on CFC Day 1 (i.e. acquisition). **c.** Representative traces of open field behavioral locomotor activity. **d.** Quantification of open field measurements to assess locomotor activity across treatment groups . **e.** Light-dark box apparatus. **f.** Quantification of time spent in each chamber of the light-dark box as an assessment of anxiety-like behavior. Analysis: One-way ANOVA, Tukey’s multiple comparisons. n = 6 animals/treatment group.

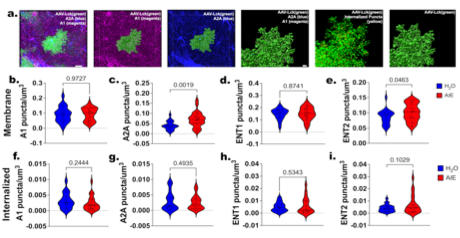
**Supplemental Fig 8. AIE induces a subtype-specific increase in adenosine receptor and transporter expression on the astrocyte membrane. a.** Representative images of astrocyte surface rendering (Lck-GFP, green), A2AR (magenta), and A1R (cyan) in the dHipp following AIE and 26-day forced abstinence period in membrane-bound and internalized. Scale bar= 5, 20 µm. **c-f**. Internalized and Membrane-bound quantification of A2A and ENT2 on the membrane of the astrocyte. (*p =*. Data are presented as violin plots representing the distribution of individual data points. Analysis: unpaired t-tests, n = 6-10 astrocytes/animal, 6-8 x 80 µm image stacks/slide, 2 slices/animal, 6 animals/treatment group.
